# A highly reproducible, safe cranial window surgery for targeted *in vivo* manipulations in the adult zebrafish telencephalon

**DOI:** 10.64898/2026.09.24.754169

**Authors:** Konstantin N. Zabegalov

## Abstract

Experimental surgery is one of the most fundamental fields of biomedical research, recreating clinical conditions on living organisms similar to clinical patients, to facilitate the design of novel therapies. For modeling neurological and psychiatric disorders, the majority of operative techniques mandatorily include craniotomy for consecutive interactions with the open brain. The most wide-spread methodology involves stereotactic surgery in rodents, implementing a specific head holder, attached to the coordinate table, providing the precise animal head positioning according to the brain structures topology. Despite the high efficiency, such systems require complex instrumental skills and long procedure duration per animal even for simple implantation protocols. Hence, novel alternative model species could be a valuable supplementary tool for such manipulations. Therefore, the zebrafish (*Danio rerio*) fits perfectly, due to its low cost, easy husbandry, rapid development, optic transparency, and high genetic and physiological similarity to rodents and humans. Furthermore, the everted zebrafish telencephalon provides excellent accessibility to critical mammalian-like structures involved in emotion processing and cognition. However, conventional avian- or rodent-inspired craniotomies in adult zebrafish often trigger inadvertent disruption of the brain vessels network, resulting in high operative mortality. A novel, geometry-driven, non-lethal cranial window protocol preserving neurovascular integrity and ensuring 90–100% post-operative survival is presented to overcome above mentioned limitations. This rapid, simple, and high reproducible technique opens new avenues for precise region-specific *in vivo* neurosurgery, high-throughput psychopharmacological screening, and neurostimulation in a powerful non-mammalian model species.

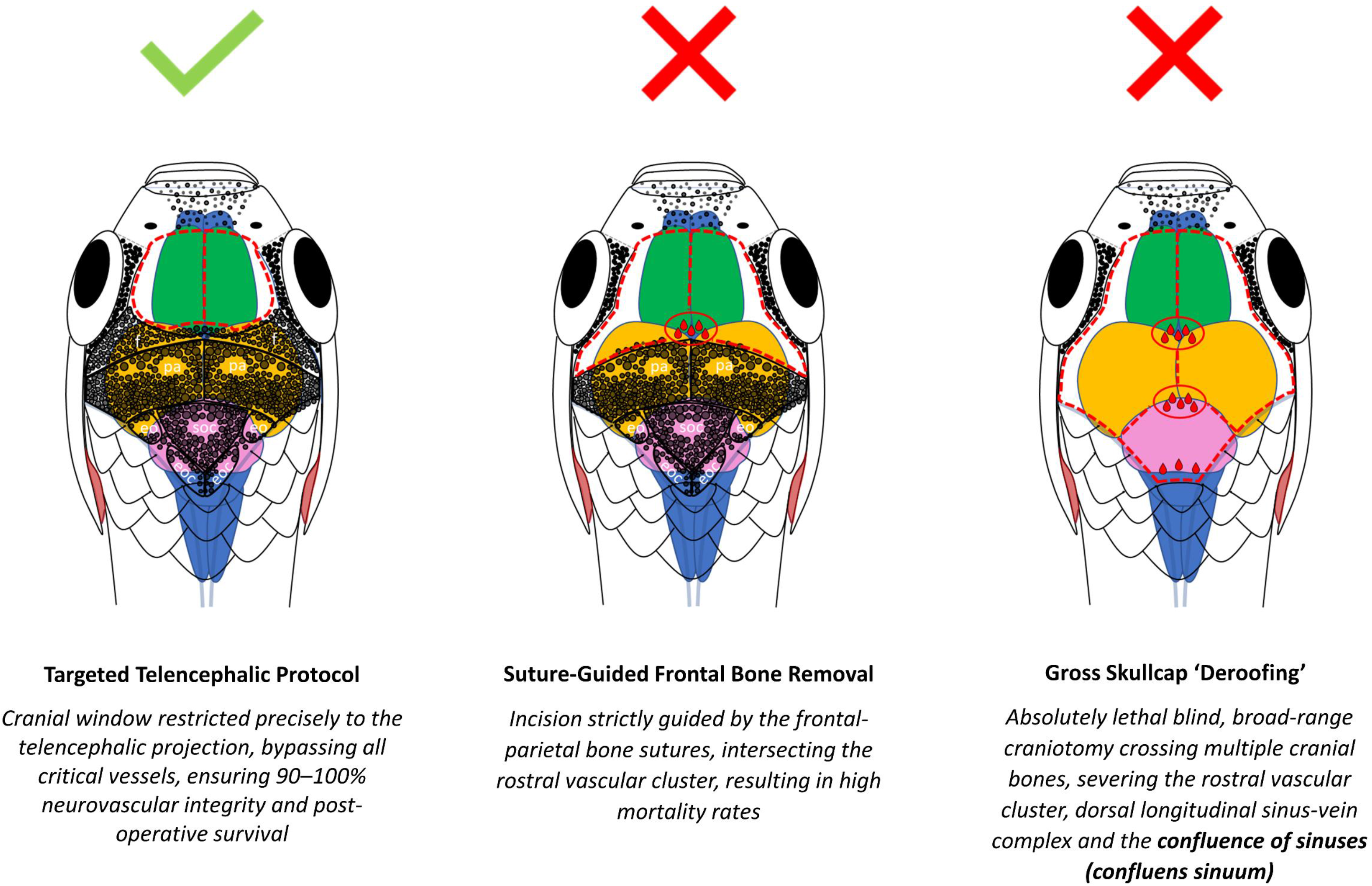

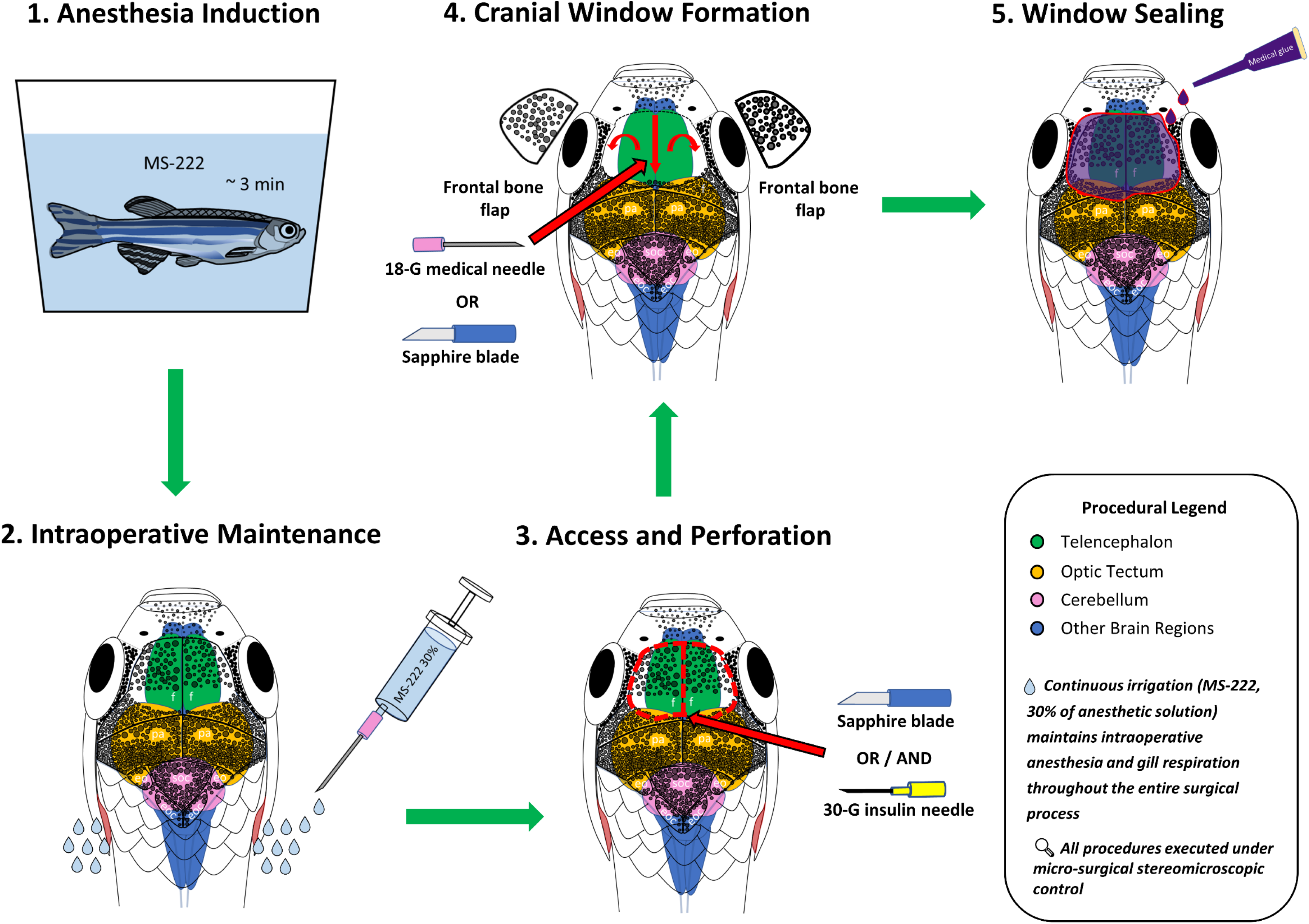

**Highlights:**

- An optimized, highly standardized craniotomy method for adult zebrafish is presented
- Targeted micro-perforation avoids major sinuses, ensuring 90–100% survival rate
- Bloodless surgery maintains functional blood-brain barrier and vascular integrity
- An automated ilastik/ImageJ pipeline provides objective trauma quantification
- The protocol bridges the gap between high-throughput screening and human neuroscience

## Introduction

Surgery as one of the major corner stones of traditional health care demands constant improvement. Corresponding procedures on laboratory animals crucially facilitate the advancement of surgical and post-operative rehabilitation protocols, biocompatible materials assessment, and implantation strategies validation, thereby providing a stable ground for the translation of these novelties into clinical practice ^1^. Although, multiple surgical simulation models are developing progressively and prone to bypass ethical restrictions, manipulations with living subjects and the dissection of post-mortem material remain the gold standard due to the unique and non-reproducible features of biological systems ^2^. As substantial evidence, 43 of 97 Nobel Prize awarded research included in vivo experiments in animal models ^3^; although smaller species have been prevailing within recent decades ^4^. In such manner, rodents (predominantly mice) are undisputed leaders among the others, possessing well-adjusted guidelines, recommendations, and legal basis for the surgical techniques ^5,6^. The general procedures mostly interfere with the cardiovascular system and heart disease models ^7–9^, the search of advanced skin and organs transplantation methods ^10,11^, orthopedic studies ^12,13^ and experimental traumatology ^14,15^, generalized cancer modeling ^16^, and the vast field of experimental neurosurgery ^17^, which is about utterly discussed in this paper.

A fundamental set up in rodent experimental neurosurgery primarily refers to stereotactic encephalometer, whose efficiency has been tested over a century from the very first application in 1908 by Horsley and Clarke ^18^. The basic principle of the device indicates 3D coordinate-based subject’s head positioning for targeted invasive (injection, implantation) or non-invasive (radiation or other optic exposures) procedures ^19–21^. As an essential supplementary, the constant improvement of brain atlases boosted the stereotactic surgery to sub-millimeter (∼ 0.5 mm) accuracy ^22,23^. Such precision made possible the design of novel sophisticated cortical trauma models, reducing the impact area to specific cortical areas (*e.g*., M1 of motor cortex and S1 of the primary somatosensory cortex) ^24^. In addition to the surgery localization, the constant refinement of implantable devices, surgery site sealing approaches and welfare scoring promotes stereotaxic manipulations to a principally new level ^25^. Eventually, these advances triggered the development of such cutting-edge therapies as the deep brain stimulation (*e.g.*, pedunculopontine and cuneiform nuclei stimulation for the gait disorders treatment ^26^), brain cancers therapies (*e.g.*, glioblastoma) with viral vectors ^27^, and viral tracing techniques applied to the deep limbic circuit studies ^28^.

Despite such impressive advances in experimental and clinical neurosurgery, rodent stereotactic procedures still remain complex for high-throughput research and clinical approaches, demanding proficient expertise in operative micromanipulations ^25,29–31^. It is widely acknowledged among experimental stereotaxic surgeons that avoiding major venous domains remains an exceptional manual challenge, heavily dependent on subjective tactile feedback. Even experienced operators frequently encounter catastrophic sinus ruptures due to slight manual over-pushing. To overcome such obstacles modern robotics actively propose novel automated flexible platforms ^32^. Although, despite their complexity and customed production, these devices have not yet reached the better time efficiency in comparison with classic manual stereotaxic surgery ^33^.

Traditionally mentioned the zebrafish (*Danio rerio*) developmental advantages, larval transparency, cost-effectiveness, easy husbandry, shared genetics and physiology with humans, simplified drug administration makes this organism an extraordinarily promising for biomedical research and high-throughput biological screening in applied fields ^34,35^. Specifically, the genetic similarity with humans indicates more than 70% general homology, more than 80% common disease-associated genes, and about 90% of the homologous genes related to the brain development disorders ^36,37^. Although the zebrafish brain possesses less sophisticated morphology and much lesser number of neurons (∼ 100, 000) than mammals, behavioral patterns they exhibit compile conservative repertoire for the most of the vertebrates (*e.g*., stress and fear responses, social preference, hunting, spatial cognition, and sleep) ^38,39^. Such conservatism is based on the general neuroanatomy plan, neurotransmission homology, and above mentioned neurogenetics, resulting in multiple clinically relevant neuropathological endophenotypes (*e.g.*, anxiety, despair, anhedonia, cognitive deficits, epilepsy, neurodevelopmental abnormalities) ^36,38,40–43^.

Regarding experimental neurosurgery, the zebrafish has a number of undisputed advantages. Primarily, the zebrafish neurocranium, constantly growing across the life-time, keeps flexibility ^44,45^, along with the lowest thickness in the “zebrafish-mouse-human” phylogenetic ladder ^46^, providing the easiest access to the brain via tiny cranial perforations, done with the typical medical needles of a micrometer gauge (*e.g.*, 30g) ^47^. Despite the current knowledge on the complex mammalian-like structure of the zebrafish meninges (double-layered dura mater and leptomeningeal barrier) ^48^, their accidental disruption does not inherently lead to catastrofic hemorrhage or lethal blood loss. The zebrafish brain vascular system possesses robust mechanisms for blood flow restoration through collateral circulation ^49^, and meningeal lymphatic-mediated revascularization ^50,51^, while the highly conserved and efficient hemostatic system rapidly controls bleeding at the injury site ^52,53^. Finally, the specific eversion (unlike mammalian evagination) of the zebrafish telencephalon ^54,55^ transposes the key basal ganglia and limbic structures from the brain depth to the very surface (*e.g.*, zebrafish hippocampus and basolateral amygdala homologues ^56,57^) ^54^, unraveling the brain *in vivo* imaging studies to the great extent.

In such manner, the novel method for the zebrafish precise telencephalic craniotomy was designed. Here, the evolution of such intrinsic methodology is shown in retrospect, tracing the historical development from early lethal skullcap ‘deroofing’ protocols (originated from the legacy methods presented in goldfish ^58,59^) compared to the final reproducible technique. Hence, the assessment of mortality rates, injury site topology, bleeding rates and the general quality of surgery process presents a wide-scaled validation, standardizing such method for the worldwide experimental neurosurgery.

## Results

### 3.1 Mortality Outcomes

The Kaplan-Meier survival analysis revealed significant differences in post-operative mortality across the four experimental sessions (Log-rank Mantel-Cox test, χ^2^ = 10.76, df = 3, *P* = 0.0131). While Session 1 demonstrated an early drop in survival, stabilizing at 43.75 % by day 7 (Figure 1 (a)), subsequent adjustments to the surgical technique led to a pronounced reduction in lethality. Crucially, a highly significant linear trend toward increased survival was observed from Session 1 to Session 4 (Log-rank test for trend, χ^2^ = 7.757, df = 1, *P* = 0.0054), with Session 4 achieving a 100% survival rate (Figure 1 (a)). This clear positive trend mathematically validates the standardization of the surgical technique and illustrates a distinct learning curve.

**Figure 1.**
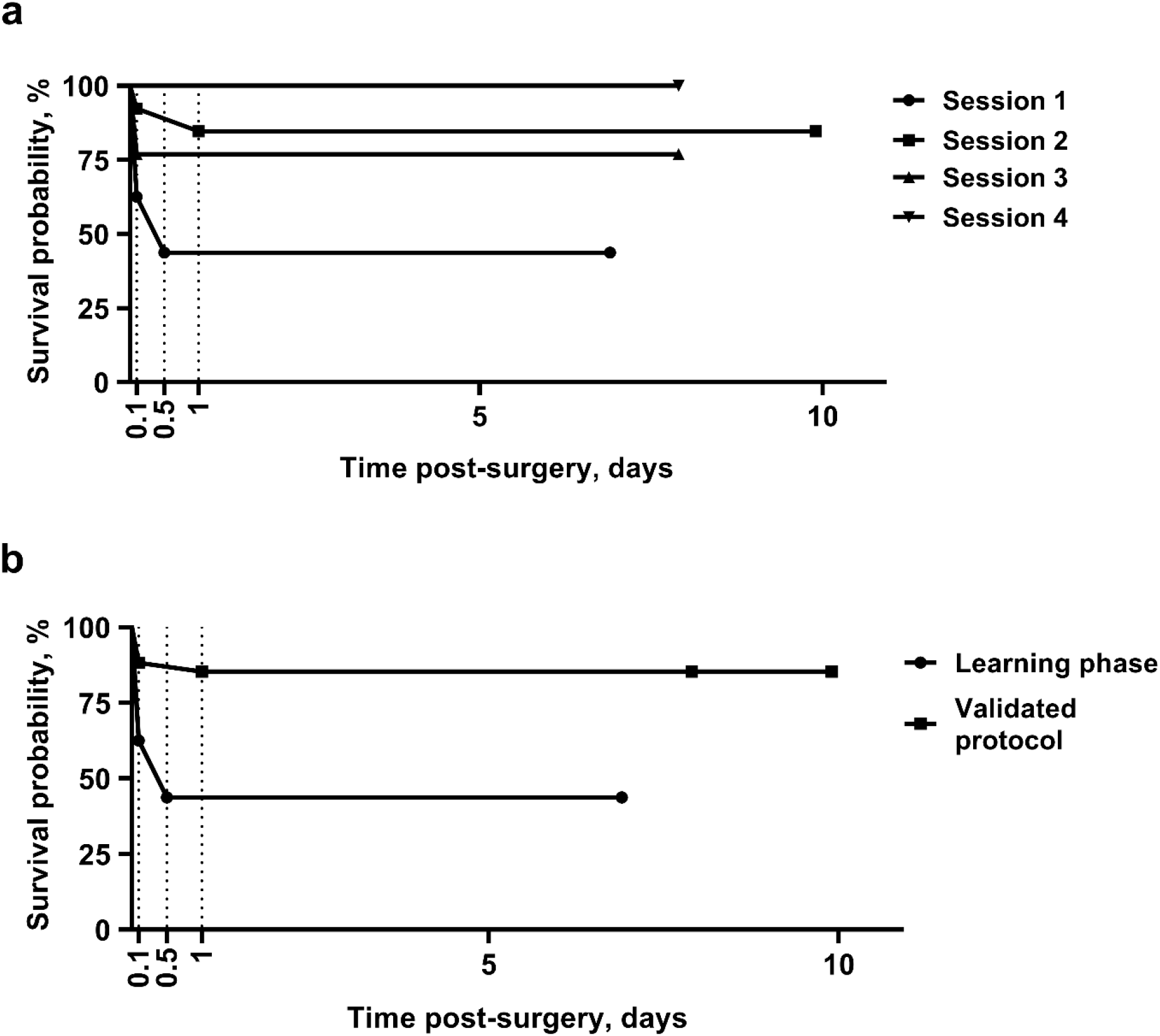
Survival proportions for Sessions 1-4 (**a**) and Learning phase including just the lethal Session 1 compared to the Validated protocol consisting of the three vital sessions (**b**) reproducing the optimized targeted telencephalic surgery. Horizontal lines on the graphs indicate the percentage of survived fish within the surgery (day 0.1), the first post-operative night (day 0.5) and the next day after the surgery (day 1) terminating with the final observation day.

Furthermore, when the data were dichotomized into two master phases (Figure 1 (b)), a highly significant increase in survival was confirmed for the validated protocol phase (Sessions 2-4 combined) compared to the initial learning phase (Session 1) (Log-rank Mantel-Cox test, χ^2^ = 9.592, df = 1, *P* = 0.0020). Strikingly, the hazard ratio (HR) calculation using the Mantel-Haenszel method demonstrated that animals in the learning phase carried a 7.19-fold higher risk of mortality than those operated under the optimized protocol (HR = 7.194, 95% CI: 2.064 to 25.08). This profound reduction in hazard firmly establishes the safety, high viability, and reproducibility of the finalized craniotomy technique.

### 3.2. Surgery Duration and Craniotomy Method Safety

The assessment of surgical efficiency within the validated phase demonstrated exceptional procedural standardization. One-way ANOVA revealed no statistically significant differences in craniotomy duration across Sessions 2,3, and 4 (*F* (2,30) = 0.4076, *P* = 0.6689) (Figure 2 (a)). This risk of statistical divergence mathematically validates the high reproducibility and consistency of the finalized ‘Targeted telencephalic protocol’. Once the baseline manual proficiency is established, the surgical execution reaches a stable plateau, ensuring predictable operation timing and minimizing extended anesthesia exposure for the animals.

**Figure 2.**
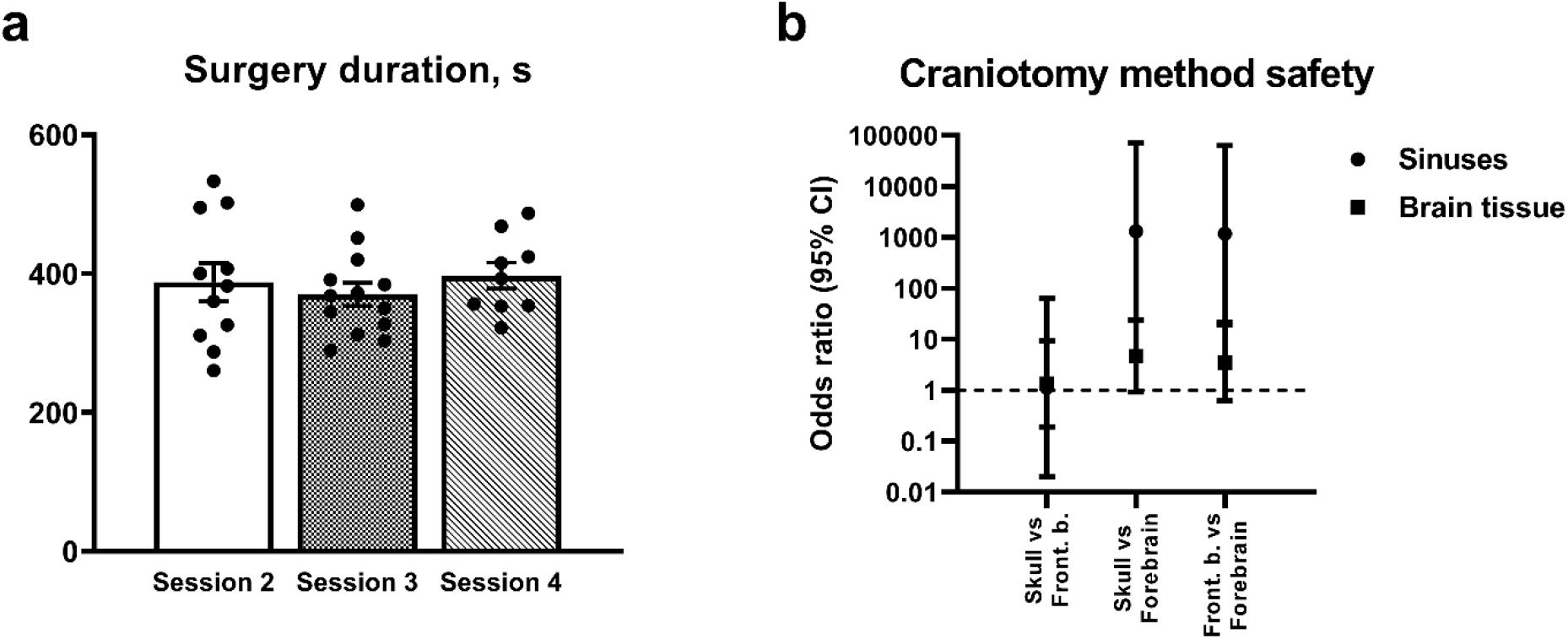
Bar chart with whiskers (**a**) presents the one-way ANOVA of time spent on the entire craniotomy process for each fish (individual dots) between sessions 2-4. Data are presented as mean with the standard error of mean (SEM) (n = 8-13). Forest plot (Y axis in the log_10_ scale) displays Odds Ratios (OR) and corresponding 95% confidence intervals (95% CI) for ‘sinuses’ (Rostral Vascular Cluster (RVC) and Dorsal Longitudinal Sinus (DLS)) disruption (circles) and parenchymal brain tissue damage (squares) across pairwise group comparisons (**b**). An OR > 1 indicates an increased risk of complications in the former group relative to the latter. Point estimates were adjusted using the Haldane-Anscombe correction (+ 0.5) to account for zero-cell frequencies. Statistical significance (*P* < 0.05) is established for venous sinus safety, as the 95% CI lower bounds do not cross the line of unity (OR = 1). Abbreviations: Skull, traditional skullcap deroofing (lethal method); Frontal b., entire frontal bone removal (lethal method); Forebrain, optimized vital telencephalic protocol.

To evaluate the comparative safety profiles of the surgical techniques, an Odds Ratio (OR) analysis with Haldane-Anscombe correction was performed across the craniotomy cohorts. The optimized vital telencephalic protocol (referred to as “Forebrain” in pairwise comparisons) demonstrated a monumental increase in vascular safety, achieving statistically significant superiority over both traditional lethal methods regarding ‘sinuses’ integrity (*P* < 0.05). Specifically, the odds of sustaining a catastrophic ‘sinuses’ disruption were 1311.00-fold higher in the skullcap group (“Skull”, OR = 1311.00, 95% CI: 24.37–70529.37) and 1173.00-fold higher in the entire frontal bone removal group (“Frontal b.”, OR = 1173.00, 95% CI: 21.67–63482.73) compared to the optimized protocol. Because the lower bounds of the 95% confidence intervals (24.37 and 21.67) remain strictly above the line of unity, the absolute safety advantage of the novel method against vascular trauma is mathematically verified (Figure 2 (b)).

In contrast, the analysis of parenchymal brain tissue damage revealed a strong clinical trend that did not reach formal statistical significance due to sample size constraints and the overlapping 95% confidence intervals (*P* > 0.05). The odds of sustaining brain tissue disruption were 4.64-fold higher in the skullcap group (95% CI: 0.92–23.48) and 3.48-fold higher in the frontal bone removal group (95% CI: 0.62–19.38) relative to the optimized forebrain protocol. Direct comparison between the two traditional lethal approaches confirmed nearly identical complication rates for both ‘sinuses’ (OR = 1.12, 95% CI: 0.02–62.74) and brain tissue integrity (OR = 1.33, 95% CI: 0.19–62.74), indicating no safety difference between the legacy methodologies (Figure 2 (b)).

### 3.3. Blood clots analysis

To verify the procedural safety and reproducibility of the optimized vital telencephalic protocol, planimetric assay of post-operative trauma and total surgery duration were compared between the exploratory learning phase (Session 1, n = 16) and the finalized validated cohort (Session 4, n = 8). According, to Shapiro-Wilk normality test, the data of all samples per each parameter analyzed were distributed normally, so the pair comparisons were processed with two-sided unpaired Student’s t test followed by Welch’s correction. Planimetric quantification of the total clot burden fraction showed no statistically significant differences between the sequential experimental phases (12.56 ± 1.467 % vs. 14.05 ± 3.150 %, respectively; unpaired t-test with Welch’s correction, t(10.14) = 0.429, *P* = 0.6767; Figure 3 (a), top). Similarly, the localized brain clots fraction (restricted to the cranial window area) remained statistically stable and unchanged across the cohorts (26.70 ± 5.484 % vs. 32.19 ± 8.809 %, respectively; unpaired t-test with Welch’s correction, t(10.29) = 0.6225, *P* = 0.5472; Figure 3 (a), middle). These findings mathematically demonstrate that the initial training period and technical familiarization do not compromise procedural safety or accelerate peripheral thrombi accumulation.

**Figure 3.**
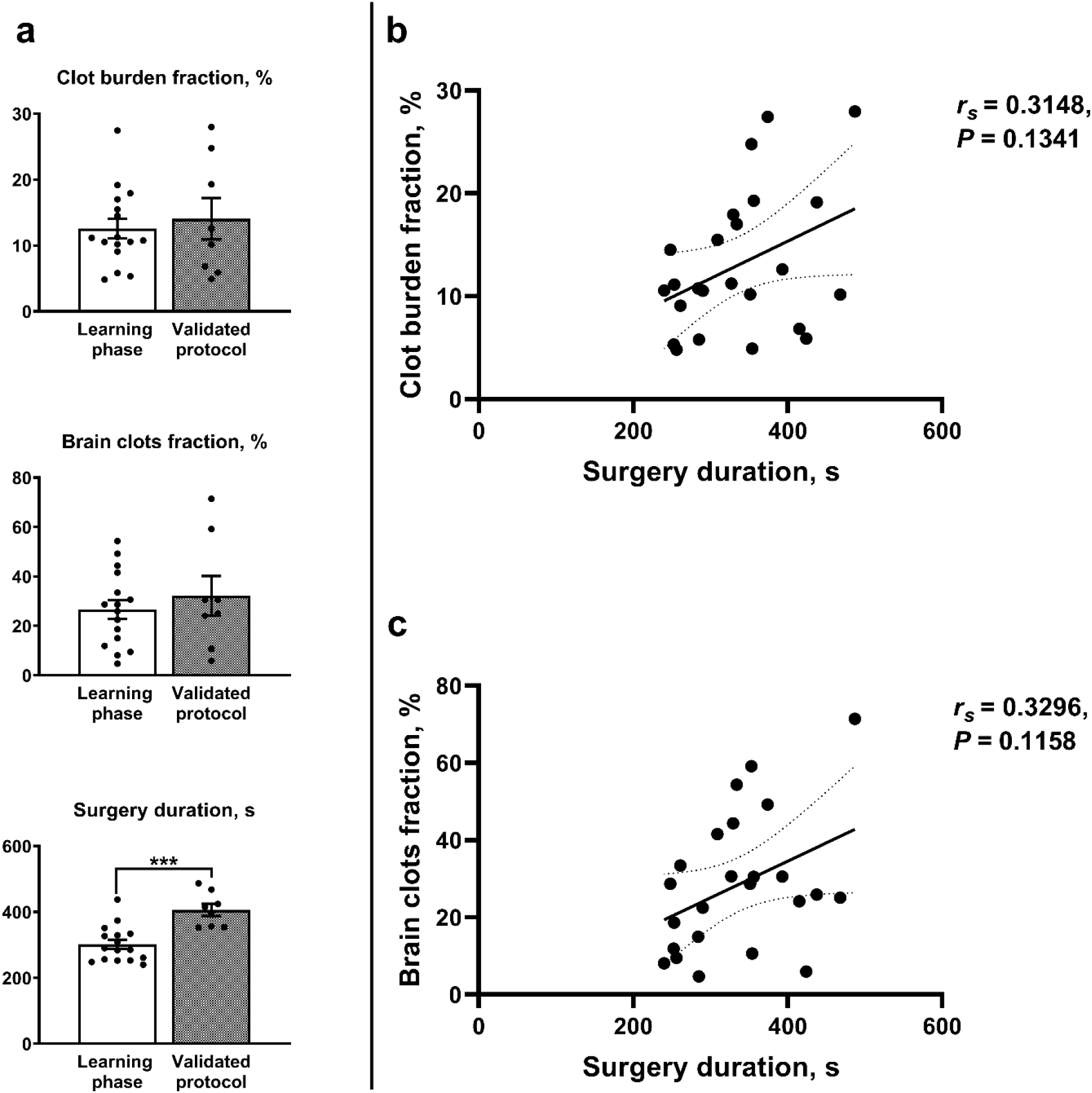
(**a**) Bar plots comparing the total clot burden fraction (top), localized brain clots fraction (middle), and total surgery duration (bottom) between the initial learning phase (n = 16) and the validated protocol cohort (n = 8). Data are presented as mean ± SEM with individual dots for each fish; *** *P* < 0.001 two-sided sunpaired Student’s t-test with Welch’s correction. (**b**, **c**) Spearman rank correlation plots evaluating the relationship between total surgery duration (seconds) and (**b**) total clot burden fraction or (**c**) localized brain clots fraction across all surgical sessions (n = 24). Solid lines indicate linear regression fits surrounded by 95% confidence bands (dotted lines). Correlation coefficients *r_s_* and corresponding two-tailed *P*-values are indicated on the plots.

However, a highly significant increase in execution time was observed upon transitioning to the validated protocol, with total surgery duration rising from 302.1 ± 104.2 seconds during the learning phase to 406.3 ± 22.92 seconds in the validation phase (302.1 ± 104.2 % vs. 406.3 ± 22.92 %, respectively; unpaired t-test with Welch’s correction, t(14.72) = 4.546, \*\*\**P* = 0.0004; Figure 3 (a), bottom). This prolonged timeline directly reflects the meticulous, high-precision micro-perforation required to safely penetrate the frontal bone along the designated cutting tracks.

To explore the potential physiological impact of extended procedural times on vascular trauma, a Spearman rank correlation analysis was performed across the combined morphometric cohort (n = 24). Crucially, the analysis revealed that the duration of the craniotomy did not actively dictate the severity of intraoperative bleeding. No statistically significant correlation was observed between surgery duration and the superficial clot burden fraction (*r_s_* = 0.3148, *P* = 0.1341; Figure 3 (b)). Similarly, the brain clots fraction showed a non-significant relationship with operational velocity (*r_s_* = 0.3296, *P* = 0.1158; Figure 3 (c)). This total lack of statistical covariance provides pivotal evidence that the localized micro-perforation technique prevents progressive vascular leakage. Even when the surgical execution required extended manual manipulation, the induced micro-trauma to the intraosseous emissary veins and the meninges remained strictly self-limiting, ensuring that prolonged procedural exposure did not exacerbate cerebral parenchymal or vascular pooling.

## Discussion

Taken into consideration measured quantitative parameters and surgical qualitative features, the targeted telencephalic cranial window protocol promotes the zebrafish craniotomy to a principally different level. Absolute vitality achieved in the Session 4 (Figure 1 (a)) makes this technique the only sustainable path to the further manipulations with the open brain, including region specific traumatization, neurophysiological registration (*e.g.*, electroencephalography / EEG, magnetoencephalography / MEG), in vivo imaging strategies (calcium imaging, two- or three-photon microscopy), micro- or nanoparticles implantation, and targeted pharmacological trials ^60–66^. Although, such historically first iterations, as the entire frontal bone removal or the gross skullcap deroofing are much faster and easier to perform, and cranial sutures serve as a reliable neuroanatomic guide for the brain structures localization (especially for an inexperienced neurosurgeon), the disruption of subjacent vascular clusters and venous sinuses leads to inevitable mortality and complete ineffectiveness for the further *in vivo* research.

Nevertheless, whether the safest targeted telencephalic surgery or the totally lethal gross-skullcap deroofing require the comprehensive map of critically important blood vessels residing the dorsal brain surface (Figure 4 (a)). According to canonical mappings of the zebrafish-related cyprinids (^67^, modified from ^68^), the major neurovascular input of the brain is provided by the arteries strictly confined to the ventromedial and intramural neuroepithelial scaffold (specifically, the midline basilar artery (BA) and its branches – central arteries (CAs)). Such localization firmly secures the brain blood supply from potentially harmful skullcap incisions and other experimental interactions with the dorsal brain surface. However, the veins on these classical maps ^67,68^ remain largely unlabeled, lacking both systematic nomenclature and depth-layer classification. Hence, the current study addresses this critical methodological and anatomical gap. By introducing a comprehensive, layer-by-layer dorsal projection of the venous drainage system (including the DLS (dorsal longitudinal sinus), RVC (rostral vascular cluster), MCeV (middle cerebral vein), and MsV (mesencephalic vein)), the first functionally annotated venous roadmap (Figure 4 (a, b)) is provided. The major basis for the neurosurgeon-oriented vascular scheme was the comprehensive classical atlas of the zebrafish vascular development by Isogai et al., 2001 ^69^, supplemented by a recent short communication on zebrafish brain vascular heterogeneity by Lee & Matsuoka, 2025 ^70^, from which all of the vessel names were adopted. Notably, while the Isogai atlas provides an invaluable segmented blueprint of the early larval (3.5 dpf) head vasculature ^69^, several critical morphological and topological divergences evolve in the adult. The dramatic volumetric expansion of the optic tectum and telencephalic hemispheres geometrically displaces the venous networks, transforming what appear as short larval bridging vessels into prominent, looping adult conduits – most notably the MsV and MCeV (Figure 4 (a, b)). This geometric displacement is heavily driven by post-embryonic developmental shifts. Recent spatiotemporal and transcriptomic atlases of the zebrafish brain vasculature confirm a pronounced transition from early, highly stereotyped lateral/superficial vascularization to extensive intraparenchymal angiogenesis and deep vessel branching during maturation ^71^. Furthermore, this structural reorganization coincides with a dramatic expansion of the neuroglial-vascular architecture, where glial coverage of intracranial vessels escalates from ∼30% in larvae to over 70% in adults ^72^. These ontogenetic expansions explain why adult drainage conduits, such as the prominent MsV and MCeV mapped in Figure 4 (b) sink into deeper anatomical planes (L2–L3) compared to their simplified, non-ossified embryonic precursors documented by Isogai et al. ^69^.

**Figure 4.**
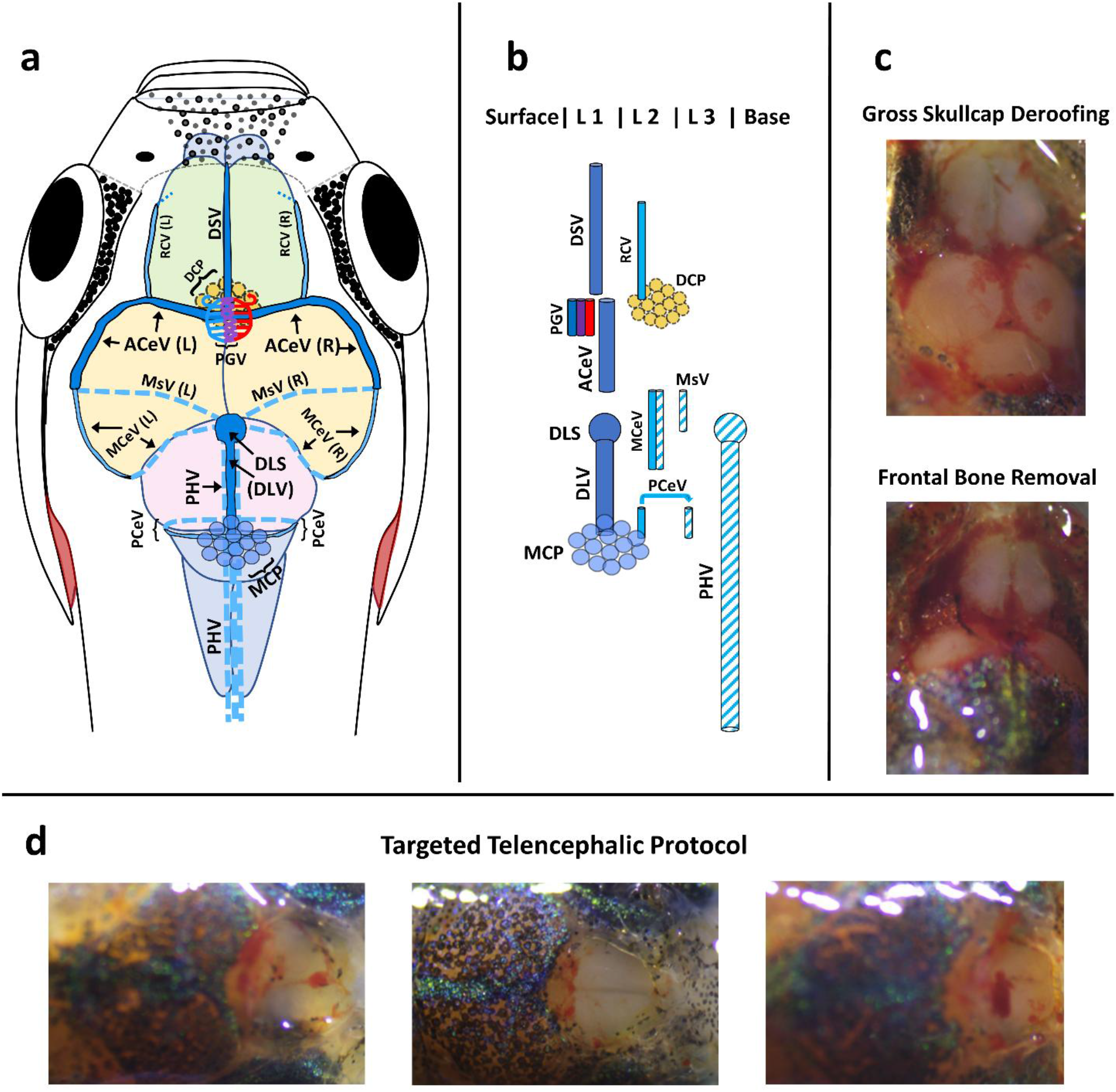
Vascular architecture of the adult zebrafish brain and craniotomy evaluation. **(a)** Dorsal view map of the open brain and cerebral vessel distribution. **(b)** Layer-by-layer (L1–L3) lateral projection of the main neurovascular and plexus domains. **(c)** Representative intraoperative photographs showing catastrophic hemorrhage following gross skullcap deroofing and frontal bone removal. **(d)** High-vitality outcomes and intact vascular integrity achieved via the targeted telencephalic protocol. The dorsal sagittal vein (DSV) tracks the frontal suture, entering the medial domain of the anterior cerebral vein (ACeV). The pineal gland vessel (PGV) forms an intricate anastomosis weaving the epiphyseal region and draining into the deeply situated diencephalic choroid plexus (DCP). Rostral cerebral veins (RCV) course along the lateral telencephalon surface, ascending toward their confluence with the lateral branches of the ACeV. The ACeV tracks the dorsolateral surface of the optic tectum, dichotomizing at the midpoint of the dorsal tectal perimeter into the mesencephalic vein (MsV) and the middle cerebral vein (MCeV). The MsV descends abruptly along the midbrain toward the neurocranial base, entering the primitive hindbrain vein (PHV) inferior to the optic tectum-cerebellum border projection. The MCeV courses along the posterior margin of the optic tectum at the second level (L2), descending along the ventral brain surface at the lateral boundary between the optic tectum and cerebellum before draining into the PHV. The leptomeningeal evagination superior to the medial boundary between the optic tectum and cerebellum forms the dorsal longitudinal sinus (DLS), which originates from the dorsal longitudinal vein (DLV) and dichotomizes into the symmetrical branches of the prosencephalic cerebral vein (PCeV) surrounding the cerebellum-crista cerebelli boundary to enter the PHV at the neurocranial basal level. A prominent and vulnerable myelencephalic choroid plexus (MCP), formed by the leptomeninx, neighbors the junction of the DLV and PCeV complexes, mirroring the vascular confluence observed during gross skullcap deroofing. While the suture-guided frontal bone removal induces severe mechanical disruption of the rostral cerebral cluster (the junction of DSV, PGV, and ACeV), RCV, and ACeV, resulting in the massive bleeding depicted in panel c (bottom), the gross skullcap deroofing (panel C; top) also pluses the critical destruction of the DLS-DLV complex and the associated posterior vascular confluence (MCP and PCeV). In contrast, even the most challenging applications of the targeted telencephalic protocol (panel d) result only in localized, minor meningeal, DSV, and/or RCV disruption, which is rapidly controlled by endogenous zebrafish hemostasis and collateral blood flow.

This layer-by-layer spatial reorganization directly dictates the functional integrity of the neurovascular unit during microsurgery. As visually validated in comparative pathological panels (Figure 4 (c, d)), legacy gross skullcap deroofing (Figure 4 (c, top)) inevitably lacerates the large, non-barricaded superficial venous conduits (specifically the DLS and the proximal stems of the MCeV, and MsV), draining enormous blood volumes into the deeply situated primitive hindbrain vein (PHV), precipitating ischemic stroke and massive hemorrhagic shock, and critically impeding IV^th^ ventricle CSF dynamics – a pathophysiological outcome mechanistically analogous to clinical reports of trapped fourth ventricle syndrome ^73^ and consistent with the zebrafish hindbrain vascular models ^67,74^. Moreover, even minor iatrogenic damage to the non-fenestrated midbrain veins (MsV and MCeV) ^70^ would obliterate the local blood-brain barrier (BBB) microenvironment, inducing uncontrolled systemic macromolecular leakage that confounds post-operative targeted pharmacological trials ^75^. Less harmful, though inadvisable, suture-guided frontal bone removal (Figure 4 (c, bottom)) falsely reassures the surgeon, yet ultimately results in fatal disruption of the ACeV (anterior cerebral vein) and PGV (pineal gland vessel) (Figure 4 (a; b, middle)), which are tightly incorporated into the leptomeningeal layer of the zebrafish complex meninx ^48,76^. Consequently, blood from the disrupted rostral vascular cluster refluxes into the MsV and MCeV, occluding midbrain venous outflow and leading to subsequent brain tissue compression (a mechanism paralleling reports in mammals on venous circuitry impairment ^77^). This often results in delayed mortality, even in fish that initially appear to have survived the procedure.

In sharp contrast, the Targeted Telencephalic Protocol (Figure 4 (d)) avoids these vascular pitfalls by navigating strictly within a localized, vessel-free bony corridor. By conserving the structural architecture of the deeper anatomical planes (L2–L3), this approach leaves the continuous, non-fenestrated capillary networks of the forebrain parenchyma fully intact ^70^, thereby preserving their physiological BBB properties. Furthermore, the microperforation trajectory successfully spares the fenestrated, highly permeable capillary clusters localized within the diencephalic and myelencephalic choroid plexuses ^70^ (DCP and MCP; Figure 4 (a)) from procedural trauma. Consequently, the bloodless cranial window established here guarantees that any subsequent therapeutic agent or molecular tracer introduced *in vivo* interacts with an uncompromised, homeostatic neural tissue, making this protocol uniquely suited for high-fidelity neuropharmacological screening and long-term *in vivo* optical imaging [2, 5].

Quantitative mathematical evaluation also favors the targeted telencephalic protocol. Hence, the conventional Kaplan-Meier survival analysis indicates a significant improvement in post-operative survival outcomes from Session 1 to Session 4. Additional sessional division into the learning phase and the validated protocol revealed a striking advantage of the novel precise telencephalic technique over classical suture-guided strategies. Furthermore, the successful standardization of the validated protocol is supported by the exceptionally consistent procedural durations across the entire surgical procedure within the Sessions 2-4, with the average duration about 6.5 minutes. However, the classical traumatic techniques are about 100 seconds shorter than the targeted telencephalic protocol, reducing the risk of an anesthesia overdose. Particularly, this time expansion is critically important for the careful telencephalon-guided frontal bone micro-perforation, providing the thousandfold minimization of the massive brain vascular disruption. However, the frontal bone micro-stabbing leads to localized meningeal and emissary veins damage, which is proved by the same blood clot burden fraction within the cranial window of the learning phase and validated protocol. Nevertheless, this micro-capillary bleeding and subsequent thrombosis never evoked detrimental life-threatening condition within the post-surgery recovery. Furthermore, the subsequent correlation analysis of the total blood clot burden and cranial window bleeding with the surgery duration completely excludes the connection between surgery prolongation and potential hemorrhagic damage.

Interestingly, there are no significant differences in the brain parenchyma damage between learning phase and validated protocol. Although, these local subjectively noted disruptions are minor, the neurogenic stem cells are presented all over the adult zebrafish brain, abundantly residing in the dorsal telencephalon ^78–80^. Experimentally tracked the entire recovery even from the deep stabbing injury of the adult zebrafish telencephalon takes about two weeks post impact ^81,82^. The dorsal telencephalon or ‘pallium’ is of particular interest for translational neuroscience, as due to developmental eversion it contains structures homologous to mammalian basal ganglia on the very surface of the brain, including medial pallium (mammalian basolateral amygdala) and lateral pallium referring to mammalian hippocampus ^54,83^. Hence, the presented protocol of telelencephalic craniotomy provides a subtle basis for the design of targeted neurotrauma or focal traumatic brain injury (TBI) models.

Despite the robust performance and high reproducibility of applied machine learning-based analysis, several methodological limitations must be acknowledged. First, the evaluation of surgical hemorrhage severity was mathematically constrained to two-dimensional (2D) stereomicroscopic projections. While 2D planimetry effectively quantifies the dorsal footprints of the clots (expressed via the Clot Burden Fraction and Brain Clots Fraction), it inherently fails to capture the three-dimensional volumetric depth, thickness, and spatial density profiles of blood accumulation. Second, this current imaging pipeline does not measure the optical density or pixel intensity within the segmented clots, making the estimation of absolute volumetric bleeding metrics mathematically challenging. Third, the specific pixel-classification boundaries are encountered when analyzing the highly lethal legacy craniotomy cohorts. In several instances of massive subdural hemorrhage, the ilastik Random Forest classifier failed to resolve ambiguous margins where the dark clots merged with the endogenous melanophore pigmentation patterns of the adult fish skullcap. Finally, the scaling is based on the approximate rostral-caudal length of the telencephalon (∼ 500 µm), disregarding the above-mentioned 3D issue. To maintain relative data validity, these complex zones were manually curated and refined using the ImageJ freehand tool. However, it is worth noting that the automated classification pipeline demonstrated high fidelity and required minimal to no manual refinement when processing cohorts subjected to the targeted forebrain protocol. The computational ambiguity between melanophores and blood clots was highly localized and primarily confined to the extracranial regions of specimens from the lethal alternative groups, where massive, unconstrained hemorrhages obscured structural boundaries. In contrast, the clean, localized nature of the non-lethal micro-perforation technique inherently minimized peripheral artifacts, thereby ensuring the robust performance of the machine-learning-based planimetric analysis without the strict necessity for phenotypic pigment-deficient models (such as *casper* or *roy*-*nacre* zebrafish lines).

Finally, it is critical to address the human factor and the intraoperative learning curve associated with the targeted telencephalic protocol. Although this methodology has been systematically engineered to be as accessible, highly standardized, and universally reproducible as possible, achieving a bloodless cranial window with an exceptionally high survival rate inherently depends on the operator’s micro-surgical experience and manual tactile feedback. Based on the cumulative validation data obtained from this study, stable and highly reproducible outcomes are typically achieved after an initial training phase of approximately 10 to 15 procedural iterations. During the first attempts, common technical pitfalls for novice operators include needle or sapphire blade inappropriate positioning or minor mechanical micro-vessel micro-trauma from over-pushing. However, because the protocol relies on strict, unambiguous skull pigmentation patterns and cranial sutures as geographical landmarks, the learning curve is significantly shorter and more predictable compared to legacy gross skullcap deroofing techniques, making this approach readily adoptable by any standard neurobiology laboratory.

In summary, a novel, high-vitality targeted craniotomy protocol for the open telencephalon of adult zebrafish has been developed. By replacing traditional, lethal skullcap deroofing and suture-guided frontal bone removal techniques with a precise, layer-by-layer micro-perforation methodology, this protocol effectively mitigates catastrophic subdural hemorrhage from major superficial venous conduits (DLS, MCeV, MsV, and ACeV). The survival rate of up to 100% achieved within the optimized validation cohort mathematically demonstrates the safety, reproducibility, and high fidelity of this approach. Furthermore, by integrating a machine learning-based planimetric pipeline, the intraoperative trauma has been successfully digitized and quantified, providing an objective framework that eliminates human observer bias. Crucially, because the protocol navigates within a localized vessel-free corridor that preserves the deeper continuous, non-fenestrated parenchymal vasculature, it leaves the blood-brain barrier functionally intact. This makes presented method uniquely suited for longitudinal *in vivo* imaging, electrophysiological mapping, and high-throughput neuropharmacological screening. Future adaptations of this baseline protocol will facilitate unprecedented, minimally invasive access to deep-seated everted brain structures, bridging the gap between high-throughput non-mammalian screening and translational human neuroscience.

## Methods

### 2.1. Animals and Housing

For this study, adult heterogenous wild-type outbred zebrafish (*Danio rerio*) were sourced from a commercial supplier (Axolotl Ltd., St. Petersburg, Russia). A total of 50 specimens were selected, maintaining an approximate 1:1 male-to-female sex ratio, with animals aged between 4 and 6 months old. The animals for surgeries were taken from the entire cohort housed together under identical baseline conditions, with individual subgroups sequentially drafted to undergo surgery across four distinct, chronologically separated sessions. The initial baseline session (Session 1, n = 16) was dedicated to evaluating alternative and legacy craniotomy approaches, specifically the highly lethal suture-guided frontal bone and gross skullcap deroofing methods. The next three consecutive sessions (Sessions 2–4, totaling n = 34 assigned fish) were performed strictly utilizing the newly developed vital telencephalic protocol to validate its safety, reproducibility, and survival outcomes. Within this validation cohort, Sessions 2 and 3 included 13 fish each. Session 4 successfully evaluated 8 operatively managed individuals; one additional animal initially allocated to this session experienced a pre-operative technical drop-out due to acute mechanical asphyxia prior to the surgical incision and was excluded from the final experimental sample size. Prior to the initiation of surgical procedures, the fish underwent a two-week acclimation period in a 60-L plastic tank equipped with continuous water filtration and aeration.

Environmental parameters were strictly regulated: water temperature was sustained at 25–27 °C, and the pH was kept within a 7.2–7.4 range. Water quality was optimized utilizing Aqualife™ conditioner (Aqualife®, Individual Entrepreneur Valkovich, Krasnodar, Russia) alongside Aquacons® antiseptic treatment (Zoomir Ltd., St. Petersburg, Russia), excluding the presence of heavy metals and reducing ammonia (NH3/NH4+), nitrite (NO2–) and nitrate (NO3–) levels to acceptable ranges (< 0.1 mg/l, < 0.3 mg/l, < 25 mg/l, respectively). Photoperiodic cycles were maintained on a stable 12-hour light / 12-hour dark regimen (lights on at 9:00 AM and off at 9:00 PM) driven by ceiling-mounted fluorescent lighting providing an intensity of 300-450 lx. Dietary management consisted of twice-daily feeding with TetraMin commercial flake food (Tetra, Ltd., Melle, Germany).

All experimental subjects were completely naïve to prior testing and underwent strict post-operative tracking, with zero exclusions during data analysis except for a single predefined lethal case that occurred independently of surgical interventions. Animal welfare, husbandry protocols, and experimental designs strictly conformed to the international zebrafish care manual ^84^, and recent FELASA-approved guidelines ^85^, Institutional Animal Care and Use Committee (IACUC) of Life Improvement by Future Technologies (LIFT) Center, LLC under the Protocol No.1, and the European Union Directive 2010/63/EU. The execution, reporting, and statistical structuring of these animal experiments strictly followed the ARRIVE guidelines, the 3Rs principles (Replacement, Reduction, and Refinement) for humane scientific research, and PREPARE guidelines. The selection of an outbred strain was dictated by population validity factors relevant to modeling central nervous system pathways. Although inbred strains offer enhanced reproducibility and genetic uniformity in neurogenetic studies, modeling complex human pathologies within genetically heterogeneous cohorts requires an outbred population approach, which offers superior translational and population validity for the goals of the current work, as well as reduced inbreeding depression ^86–88^.

### 2.2. Surgical Protocol and Cranial Window Formatting

Surgical procedures were executed across four distinct chronological sessions as detailed in Section 2.1. During the initial session, alternative and legacy methods (frontal bone and gross skullcap deroofing) were performed to establish comparative baseline lethality. The subsequent three validation sessions were carried out strictly according to the optimized vital protocol described below (see Supplementary Video 1).

#### 2.2.1. Anesthesia Administration and Maintenance

An adult wild-type heterogeneous outbred zebrafish (*Danio rerio*) was transferred from the holding facility to a plastic anesthesia exposure tank containing 300 mL of tricaine methanesulfonate (MS-222, 150 mg/L; Merck Group, Darmstadt, Germany). To prevent metabolic acidosis, the solution was buffered with sodium bicarbonate (NaHCO3, conforming to GOST 2156-76 standard for food and pharmaceutical chemical parameters) at a strict 1:2 ratio following validated guidelines ^89^. The animal was monitored until it reached the deep surgical anesthetic plane, characterized by the complete absence of the tail-pinch reflex and a profound reduction in opercular movements (usually within 3 min). During the brief transfer and positioning phase, a minimal residual caudal reflex was tolerated only prior to the initiation of the surgical maintenance protocol, provided that consistent surgical plane anesthesia was immediately re-established and sustained throughout the subsequent craniotomy using the irrigation system described below.

#### 2.2.2. Animal Positioning and Intraoperative Maintenance

Upon reaching the targeted anesthetic plane, the fish was extracted from the anesthesia exposure tank and positioned ventrally downward (dorsal side upward) in its natural anatomical posture on a customized surgical platform. The platform featured a contoured shallow groove (0.3 – 1 mm height) matching the fish’s body morphology, bounded laterally by the heads of office pins to prevent displacement while strictly avoiding mechanical pressure on the opercula and gills. To maintain physiological temperature and mitigate post-operative shock, the entire apparatus was placed over a napkin-wrapped cooling agent on a stereomicroscope stage under a trinocular stereomicroscope (NSZ608T, Novel Optics, Ningbo, China). Surgical anesthesia, immobilization, and proper opercular/cutaneous respiration were continuously maintained by regularly irrigating the gills with a maintenance anesthetic solution (45 mg/L MS-222, representing 30% of the initial induction concentration), ensuring the complete absence of any nociceptive or movement responses during the entire micro-incision process.

#### 2.2.3. Craniotomy and Telencephalic Window Formation

The caudal aspect of the head was gently stabilized using ophthalmic micro-forceps with pin-shaped branches. Initial micro-perforations were executed along the perimeter of the telencephalic projection at a 45° angle relative to the frontal bone surface. These precise incisions were performed using either a 30-G insulin needle or a 0.2-mm thick, 30° single-edge or double-edge sapphire blade. Subsequently, the dashed micro-perforations in the frontal bone were joined into a continuous, consolidated osteotomy line using the cutting edge of an 18-G medical needle or the previously mentioned sapphire blade. To ensure maximum cutting efficiency, both the 30-G and 18-G needles were replaced after each individual surgery. While the 18-G needle demonstrated superior durability for routine bone reflections, the sapphire blade provided optimal precision, despite its high fragility requiring replacement every 4 to 5 procedures. The final craniotomy phase involved the lateral reflection of the fronto-parietal bone segment overlying the telencephalon. The bone flap was flipped laterally in a hinge-like manner from the metopic suture toward the lateral margin of the cranium using the 18-G needle or sapphire blade. The absence of acute nociceptive responses was further validated by a stable and rhythmic opercular respiration rate, which consistently maintained a mean baseline of approximately 120 opercular movements per 30 seconds throughout the post-craniotomy windowing phase (as documented in Supplementary Video 1). No tachyarrhythmia or abrupt respiratory arrests were observed during bone manipulation.

#### 2.2.4. Post-Surgical Debridement and Cranial Sealing

To prevent cerebral contamination, any inadvertent ingress of the anesthetic solution onto the cranial surface or into the window was mitigated by removing residual fluid with a clean, fiber-free napkin (Kimwipes®; Kimberly-Clark Corp., Irving, TX, USA) or the fiber-less side of a clean multi-purpose laboratory tissue (Supplementary video 1, ‘Draining the Surgery Site’). To minimize the risk of bacterial contamination, the tank water and anesthetic vehicle were enriched with an antiseptic additive (Aquacons®; Zoomir Ltd., St. Petersburg, Russia). Following the designated experimental intervention, if the reflected bone flap remained structurally intact and unfractured, it was gently repositioned back into its native anatomical site following its original trajectory to ensure precise closure of the cranial window. In some cases, the backside positioning of the bone flap is acceptable if it maximally simulates the initial structure (Supplementary video 1). If bone fragmentation occurred during the reflection or removal process, a meticulous piece-by-piece skullcap restoration was performed. The final step of the protocol involved hermetic sealing of the surgical site using a biocompatible medical cyanoacrylate glue (Histoacryl®; B. Braun Melsungen SE, Germany). The adhesive application was initiated strictly at the triple border junction between the two operated telencephalic lobes and the remaining caudal frontal bone. The glue was delivered in micro-droplets and gently spread across the craniotomy perimeter to secure unstable bone fragments and ensure full structural integrity.

### 2.3. Mortality and Survival Analysis

Post-operative survival and mortality profiles were systematically monitored across all cohorts. To accurately capture the kinetics of acute and delayed mortality, specialized close-interval tracking was implemented within the first 24 hours post-surgery. Specifically, survival status was recorded at three critical intervals: immediately after the procedure to log intraoperative mortality (designated as day 0.1), following the first post-operative night (designated as day 0.5), and on the subsequent day (day 1). Thereafter, monitoring was performed daily until the pre-established experimental endpoints of each chronological cohort were reached (7 days for Session 1, 10 days for Session 2, and 8 days for Sessions 3 and 4). Intraoperative cardiac or respiratory arrest, or a prolonged absence of opercular movements exceeding 5 minutes during or immediately after the procedure, were classified as acute procedural mortality.

For long-term validation, cumulative survival probabilities were calculated using the Kaplan-Meier estimator, mathematically accommodating the variable tracking windows via standard right-censoring. Statistical comparisons of survival kinetics were performed using the two-sided log-rank (Mantel-Cox) test. Global operational variability across all four chronological sessions was assessed first, supplemented by the two-sided log-rank test for trend to evaluate directional improvements in procedural safety over time. To explicitly quantify the surgical efficacy of the newly developed method, a focused log-rank comparison was conducted between the initial exploratory learning phase (Session 1, legacy protocols) and the pooled validation cohort managed under the optimized vital telencephalic protocol (Sessions 2–4). The relative mortality risk was quantified using the Mantel-Haenszel hazard ratio (HR) calculation. All statistical models and curve fittings were generated using GraphPad Prism software (v.9.3.1, GraphPad Software, San Diego, CA, USA), maintaining a two-tailed *P*-value of less than 0.05 as the threshold for statistical significance.

### 2.4. Surgical Safety, Duration, and Complication Metrics

Intraoperative safety profiles, operational duration, and acute procedural complication rates were systematically evaluated across all experimental groups. The occurrence of major surgical complications – specifically catastrophic intracranial hemorrhage from rostral vascular cluster (RVC), including the conflux of dorsal sagittal vein (DSV) and pineal gland vessel (PGV), or dorsal longitudinal sinus (DLS) disruption (labeled in the result section as ‘sinuses’), accidental parenchymal tissue damage, and acute respiratory failure—was recorded as a binary outcome (presence or absence of complication per animal). Severe bleeding was operationalized as any visible blood accumulation within the cranial window space immediately following bone reflection. To evaluate the technical reproducibility and standardization of the vital protocol, total surgery duration was precisely clocked in seconds, spanning from the initial perforation of the frontal bone to the complete polymerization and setting of the cyanoacrylate matrix sealing the cranial window. Differences in surgery duration across the validation cohorts (Sessions 2–4) were statistically evaluated using ordinary two-sided one-way analysis of variance (ANOVA) followed by Tukey’s post-hoc test for multiple comparisons.

To statistically quantify the relative safety and clinical odds of experiencing a procedural complication under different craniotomy approaches, a comprehensive Odds Ratio (OR) analysis was performed. The legacy protocols (suture-guided frontal bone and gross skullcap deroofing methods) were treated as the baseline exposure cohorts to evaluate the risk reduction achieved by the optimized vital telencephalic protocol. Due to the presence of zero-cell frequencies in the contingency tables for specific complication categories, the Haldane-Anscombe correction was applied by adding 0.5 to all cells to eliminate mathematical indeterminacy, stabilize the log-odds standard error, and prevent division-by-zero errors. Point estimates for the Odds Ratios along with their corresponding 95% confidence intervals (95% CI) were calculated using the standard error of the log-odds method. The resulting parameters were systematically structured to generate a Forest Plot visualization to illustrate comparative procedural risk profiles. All statistical computations and graphical models were executed using Microsoft Excel and GraphPad Prism software (v.9.3.1, GraphPad Software, San Diego, CA, USA), maintaining a strict significance threshold of *P* < 0.05.

### 2.5. Cerebral Micro-Hematoma and Blood Clots Analysis

To quantitatively evaluate post-operative vascular integrity and parenchymal trauma without human observer bias, an automated, machine learning-based planimetric pipeline was developed. Following bone reflection, high-resolution digital screenshots of the exposed cranial window were taken of the surgery videos captured with the stereomicroscope (NSZ608T, Novel Optics, Ningbo, China) equipped with a digital CMOS camera (ToupCam E3ISPM08300KPC, ToupTek Photonics Co., Ltd., Hangzhou, China) under standardized stereomicroscope illumination kept constant for the analyzed cohorts. Due to subtle variations in camera zoom and minor fish positioning tilt, spatial normalization was accomplished by utilizing the standard rostro-caudal length of the adult zebrafish telencephalon (∼500 µm) as an internal biological calibration standard for each specimen.

Semi-automated image segmentation was conducted using a coupled ilastik software (v1.4.0, ilastik Team, European Molecular Biology Laboratory, Heidelberg, Germany) and ImageJ software (v1.54, National Institutes of Health, Bethesda, MD, USA) workflow. First, broad anatomical regions of interest (ROIs) were manually outlined in ImageJ to define the ***Cranial Window Area*** (restricted by the perimeter of the physical bone cut) and the ***Blood Clots Residence Area*** (the entire head surface site, containing blood clots resulted from the surgery). To isolate blood clots (the most intensively colored spots), raw RGB screenshots were processed in ilastik utilizing a Random Forest pixel classifier. The pixel classification workflow was interactively trained to differentiate two distinct labels (Clot vs. Background) based on color/intensity, edge, and texture features across multiple scale sizes (σ = 0.3 to σ = 5.0). The resulting predictions were exported as 8-bit binary segmentation masks. Due to inherent limitations of two-dimensional stereomicroscopy, which complicates automated volumetric/intensity estimation of blood accumulation and may cause heavy hemorrhage to visually blend with dense melanophore pigmentation patterns, all machine-learning masks underwent mandatory expert visual quality control. Any missed or overlapping clot perimeters, particularly within the highly lethal legacy protocol cohorts, were manually corrected and refined using the ImageJ freehand tool (marked as unnamed areas in the ROI manager archive-files) before being locked into the final ROI files to ensure maximum data validity.

These binary masks were subsequently imported into ImageJ, thresholded, and loaded into the ROI Manager. To mathematically restrict calculations to specific tissue surfaces, logical ‘AND’ operations were executed within the math engine of the ROI Manager to extract exact pixel intersections. Two definitive normalized metrics were calculated: the Clot Burden Fraction (the percentage of total clot area within the Blood Clots Residence Area) and the Brain Clots Fraction (the percentage of localized clot area restricted to the Cranial Window Area).

Statistical comparisons of these planimetric fractions and total surgery duration were conducted between the exploratory learning phase (Session 1, legacy protocols) and the final optimized reference cohort (Session 4, validated protocol) using either a two-tailed unpaired Student’s t-test with Welch’s correction or a two-tailed non-parametric Mann-Whitney U test, depending on data normality (checked with Shapiro-Wilk test). This focused two-cohort comparison was implemented to explicitly contrast the physiological extremes of procedural trauma and highlight the maximum achievable optimization threshold. To evaluate the relationship between procedural pacing and bleeding severity within these contrasting cohorts, two-sided non-parametric Spearman rank correlation coefficients (*rs*) were computed between total surgery duration (seconds) and both planimetric clot fractions. Statistical modeling and regression lines with 95% confidence bands were generated using GraphPad Prism software (GraphPad Software, San Diego, CA, USA), with significance maintained at *p* < 0.05.

### 2.6. Experiment termination and euthanasia

Upon completion of the post-surgical monitoring period, all zebrafish (Danio rerio) were humanely euthanized via gradual cooling (hypothermic shock) in accordance with the protocol described by Collymore et al. (2014). Briefly, fish were transferred to a dedicated euthanasia chamber filled with fresh filtered system water. The chamber was placed within a larger insulated container, and crushed ice was progressively added to the outer container to decrease the water temperature in the inner chamber at a rate of approximately 1°C per minute until it reached 4°C. The fish were maintained at this temperature, and the loss of opercular movement and lack of response to tactile stimuli were monitored for at least 20 minutes to confirm death. Following euthanasia, a subset of brain specimens was tissue-fixed in 4% paraformaldehyde (PFA) for retrospective morphological archiving, while the remaining biological waste was strictly disposed of as Class A (practically non-hazardous) material in compliance with the relevant veterinary regulations and national biosafety guidelines (Article 4.3 of the Law of the Russian Federation No. 4979-1 “On Veterinary Medicine” (dated May 14, 1993)).

## Declaration of Generative AI and AI-assisted Technologies in the Writing Process

The sole author used large language models (LLMs) during the preparation of this manuscript strictly for stylistic refining, grammatical correction, and linguistic optimization of the text. After using this tool, the author thoroughly reviewed and edited the content to ensure scientific accuracy, and takes full responsibility for the integrity of the data, interpretations, and the final layout of the manuscript.

## Acknowledgments

The author sincerely thanks the Life Improvement by Future Technologies (LIFT) Center, LLC for providing the infrastructure, micro-imaging facilities, and specialized stereomicroscopic camera equipment that made this study possible.

## Author Contributions

The sole author KNZ conceived and designed the study; developed the surgical protocol and performed all microsurgical procedures; designed and trained the machine-learning pipeline for planimetric analysis; performed all statistical processing, conceptualized and drew the anatomical vascular mapping, wrote the manuscript, and curated the open-access data repository.

## Ethics approval and animal welfare declaration

All experimental protocols, animal husbandry procedures, and surgical interventions involving adult zebrafish (*Danio rerio*) in this study were performed in strict accordance with international bioethical standards, including the principles of the 3Rs, the ARRIVE guidelines, and the PREPARE guidelines. To ensure rigorous ethical compliance, the specific surgical and experimental protocols for this study were formally reviewed, authorized, and approved by the Research Ethics Committee (REC) of Life Improvement by Future Technologies (LIFT) Center via the Protocol No.1 on November 14, 2025 (statement issued on November 17, 2025). All surgical procedures were conducted under the direct, independent oversight of a designated animal welfare officer to minimize any potential pain or distress, and to ensure strict adherence to the authorized protocol (see Supplementary Video 1).

## Data Availability

The high-resolution, step-by-step surgical video (Supplementary Video 1) is permanently archived and publicly accessible via Figshare at https://figshare.com/s/03223d34ae7b66262695. Pixel segmentation, classification, and planimetric analyses were executed using publicly available open-source software platforms, specifically ilastik (v1.4.0) and ImageJ (Fiji), utilizing the standard built-in Random Forest workflow parameters as described in the Methods section. No custom software code or proprietary algorithms were developed for this study. The trained ilastik classifier projects used during the validation phase are maintained under institutional archiving and are available to editors and reviewers upon request to the corresponding author.

**Supplementary Video 1.** Step-by-step micro-surgical execution of the stabilized cranial window surgery in an adult wild-type zebrafish, demonstrating animal immobilization, continuous opercular irrigation, skull micro-perforation, and hermetic sealing. High-resolution file access is detailed in the Data Availability statement.

## References

1 Choudhary, O. P. Animal models for surgeries and implants: a vital tool in medical research and development. Annals of Medicine and Surgery 87 (2025).

2 Davidson, E. L., Penniston, K. L. & Farhat, W. A. Advancements in surgical education: exploring animal and simulation models in fetal and neonatal surgery training. Frontiers in Pediatrics Volume 12 - 2024, doi:10.3389/fped.2024.1402596 (2024).

3 Hubert, T. Advocacy for Adequate Translational Surgery in Large Mammals. European surgical research. Europaische chirurgische Forschung. Recherches chirurgicales europeennes 66, 46–48, doi:10.1159/000546174 (2025).

4 Society, U. A. R. Nobel Prizes (Physiology and Medicine), <https://www.animalresearch.info/en/medical-advances/nobel-prizes/> (2025).

5 Gantenbein, F. et al. Protocol for a systematic review of good surgical practice guidelines for experimental rodent surgery. BMJ open science 6, e100280, doi:10.1136/bmjos-2022-100280 (2022).

6 National Research Council Committee for the Update of the Guide for the, C. & Use of Laboratory, A. in Guide for the Care and Use of Laboratory Animals (National Academies Press (US) Copyright © 2011, National Academy of Sciences., 2011).

7 Farag, A. et al. A review on experimental surgical models and anesthetic protocols of heart failure in rats. Frontiers in Veterinary Science Volume 10 - 2023, doi:10.3389/fvets.2023.1103229 (2023).

8 Holt, A. W. & Tulis, D. A. Experimental Rat and Mouse Carotid Artery Surgery: Injury & Remodeling Studies. ISRN minimally invasive surgery 2013, doi:10.1155/2013/167407 (2013).

9 Jiang, Y. et al. Protocol for renal artery embolization via caudal artery in rats. STAR Protocols 6, 104080, 10.1016/j.xpro.2025.104080 (2025).

10 Schmauss, D., Weinzierl, A., Schmauss, V. & Harder, Y. Common Rodent Flap Models in Experimental Surgery. European surgical research. Europaische chirurgische Forschung. Recherches chirurgicales europeennes 59, 255–264, doi:10.1159/000492414 (2018).

11 Wenzel, N., Blasczyk, R. & Figueiredo, C. Animal Models in Allogenic Solid Organ Transplantation. Transplantology 2, 412–424 (2021).

12 Chen, F., Schiffer, N. E. & Song, J. Animal Models of Orthopedic Implant-Associated Infections and Revisions. ACS Biomaterials Science & Engineering 11, 2052–2068, doi:10.1021/acsbiomaterials.4c02331 (2025).

13 Pan, Y. & Cohen, S. Reporting practices of anesthetic and analgesic use in rodent orthopedic research. Scientific reports 14, 26225, doi:10.1038/s41598-024-76750-x (2024).

14 Lang, A., Schulz, A., Ellinghaus, A. & Schmidt-Bleek, K. Osteotomy models - the current status on pain scoring and management in small rodents. Laboratory animals 50, 433–441, doi:10.1177/0023677216675007 (2016).

15 Ponz-Lueza, V., Lopiz, Y., Arvinius, C., Rodriguez-Bobada, C. & Marco, F. in Animal Models in Medical Research (ed Pınar Atukeren) (IntechOpen, 2024).

16 De Vleeschauwer, S. I. et al. OBSERVE: guidelines for the refinement of rodent cancer models. Nature Protocols 19, 2571–2596, doi:10.1038/s41596-024-00998-w (2024).

17 Janowski, M. Experimental Neurosurgery in Animal Models. (Humana New York, NY, 2016).

18 Horsley, V. & Clarke, R. H. THE STRUCTURE AND FUNCTIONS OF THE CEREBELLUM EXAMINED BY A NEW METHOD. Brain 31, 45–124, doi:10.1093/brain/31.1.45 (1908).

19 Assi, H., Candolfi, M., Lowenstein, P. R. & Castro, M. G. Rodent Glioma Models: Intracranial Stereotactic Allografts and Xenografts. Neuromethods 77, 229–243, doi:10.1007/7657_2011_33 (2012).

20 Cecyn, M. N. & Abrahao, K. P. Where do you measure the Bregma for rodent stereotaxic surgery? IBRO Neuroscience Reports 15, 143–148, 10.1016/j.ibneur.2023.07.003 (2023).

21 Geiger, B. M., Frank, L. E., Caldera-Siu, A. D. & Pothos, E. N. Survivable stereotaxic surgery in rodents. Journal of visualized experiments : JoVE, doi:10.3791/880 (2008).

22 Conti, A. et al. A Brief History of Stereotactic Atlases: Their Evolution and Importance in Stereotactic Neurosurgery. Brain sciences 13, doi:10.3390/brainsci13050830 (2023).

23 Spyrantis, A. et al. Accuracy of Robotic and Frame-Based Stereotactic Neurosurgery in a Phantom Model. Frontiers in Neurorobotics Volume 16 - 2022, doi:10.3389/fnbot.2022.762317 (2022).

24 Wechakarn, P. et al. Modified stereotactic neurosurgery techniques for rodent surgery enhance survival and reduce surgery time in a severe traumatic brain injury model. Scientific reports 15, 22166, doi:10.1038/s41598-025-05328-y (2025).

25 Pérez-Martín, E. et al. Refining Stereotaxic Neurosurgery Techniques and Welfare Assessment for Long-Term Intracerebroventricular Device Implantation in Rodents. Animals 13, 2627 (2023).

26 Hofer, A. S. et al. Stereotactic rodent-to-human approximation of the mesencephalic cuneiform nucleus to guide deep brain stimulation. Brain stimulation 19, 103066, doi:10.1016/j.brs.2026.103066 (2026).

27 Chakraborty, S. et al. Triple-Survival Stereotactic Brain Surgeries for the Intracranial Injections of Glioblastoma Stem-like Cells and Oncolytic Herpes Simplex Viruses. Methods and Protocols 9, 82 (2026).

28 Wang, Q. et al. Genoarchitecture and input–output organization of the mouse basal ganglia and thalamic parafascicular nucleus. Nature Neuroscience 29, 1248–1264, doi:10.1038/s41593-026-02253-9 (2026).

29 Azimzadeh, M., Mohd Azmi, M. A. N., Reisi, P., Cheah, P. S. & Ling, K. H. Step-by-step approach: Stereotaxic surgery for in vivo extracellular field potential recording at the rat Schaffer collateral-CA1 synapse using the eLab system. MethodsX 12, 102544, doi:10.1016/j.mex.2023.102544 (2024).

30 Ferry, B. & Gervasoni, D. Improving Stereotaxic Neurosurgery Techniques and Procedures Greatly Reduces the Number of Rats Used per Experimental Group-A Practice Report. Animals : an open access journal from MDPI 11, doi:10.3390/ani11092662 (2021).

31 Simonian, A., Obeid, G. & Cusack, L. Surgical correction of unilateral entropion in a Syrian hamster. Journal of Exotic Pet Medicine 55, 28–31, 10.1053/j.jepm.2025.09.001 (2025).

32 Ly, P. T. et al. Robotic stereotaxic system based on 3D skull reconstruction to improve surgical accuracy and speed. Journal of neuroscience methods 347, 108955, doi:10.1016/j.jneumeth.2020.108955 (2021).

33 Machetanz, K. et al. Time Efficiency in Stereotactic Robot-Assisted Surgery: An Appraisal of the Surgical Procedure and Surgeon’s Learning Curve. Stereotactic and Functional Neurosurgery 99, 25–33, doi:10.1159/000510107 (2020).

34 Podlasz, P., Migocka-Patrzałek, M., Prajsnar, T. K., Sarosiak, A. & Tylzanowski, P. Recommendations of the Polish Zebrafish Society on the use of the zebrafish (Danio rerio) model in biomedical research. Acta Biochimica Polonica Volume 73 - 2026, doi:10.3389/abp.2026.16545 (2026).

35 Siddiqui, S., Siddiqui, H., Riguene, E. & Nomikos, M. Zebrafish: A Versatile and Powerful Model for Biomedical Research. BioEssays 47, e70080, 10.1002/bies.70080 (2025).

36 Burgess, H. A. & Burton, E. A. A Critical Review of Zebrafish Neurological Disease Models−1. The Premise: Neuroanatomical, Cellular and Genetic Homology and Experimental Tractability. Oxford Open Neuroscience 2, kvac018, doi:10.1093/oons/kvac018 (2023).

37 Howe, K. et al. The zebrafish reference genome sequence and its relationship to the human genome. Nature 496, 498–503, doi:10.1038/nature12111 (2013).

38 Pang, M. et al. in Zebrafish Model in Medical Research (ed Geonildo Rodrigo Disner) (IntechOpen, 2025).

39 Randlett, O. et al. Whole-brain activity mapping onto a zebrafish brain atlas. Nature methods 12, 1039–1046, doi:10.1038/nmeth.3581 (2015).

40 da Silva Junior, F. C., de Sousa, I. P., Mamede, J. P. M., Rosemberg, D. B. & Luchiari, A. The anoxia escape test as a novel and sensitive protocol to assess despair-like behavior in zebrafish. Journal of neuroscience methods 432, 110775, doi:10.1016/j.jneumeth.2026.110775 (2026).

41 Pansera, L. et al. Zebrafish as an Integrative Model for Central Nervous System Research: Current Advances and Translational Perspectives. Life 15, 1751 (2025).

42 Prakash, K. A., Nagmoti, D. S., Borkar, M. S., Pannalal, H. K. & Bandaru, N. Review on zebra fish as an alternative animal model for neurological studies. Advances in Biomarker Sciences and Technology 7, 320–334, 10.1016/j.abst.2025.08.005 (2025).

43 Wang, L. et al. Advances in Zebrafish as a Comprehensive Model of Mental Disorders. Depression and Anxiety 2023, 6663141, 10.1155/2023/6663141 (2023).

44 Mork, L. & Crump, G. Zebrafish Craniofacial Development: A Window into Early Patterning. Current topics in developmental biology 115, 235–269, doi:10.1016/bs.ctdb.2015.07.001 (2015).

45 Topczewska, J. M., Shoela, R. A., Tomaszewski, J. P., Mirmira, R. B. & Gosain, A. K. The Morphogenesis of Cranial Sutures in Zebrafish. PLOS ONE 11, e0165775, doi:10.1371/journal.pone.0165775 (2016).

46 Liao, Y. et al. In vivo quantitative evaluation of the relationship between skull thickness and body length and age in zebrafish using OCT. Authorea 2023, doi:doi:10.22541/au.168733858.87144720/v1 (2023).

47 Kizil, C., Iltzsche, A., Kaslin, J. & Brand, M. Micromanipulation of gene expression in the adult zebrafish brain using cerebroventricular microinjection of morpholino oligonucleotides. Journal of visualized experiments : JoVE, e50415, doi:10.3791/50415 (2013).

48 Galanternik, M. V., et al. Anatomical and Molecular Characterization of the Zebrafish Meninges. bioRxiv : the preprint server for biology, doi:10.1101/2025.04.09.646894 (2025).

49 Xia, I. F., et al. A mitochondrial program encodes brain vascular reserve. (2026).

50 Chen, J. et al. Acute brain vascular regeneration occurs via lymphatic transdifferentiation. Developmental cell 56, 3115–3127.e3116, doi:10.1016/j.devcel.2021.09.005 (2021).

51 Kraus, A., et al. Live longitudinal imaging of meningeal cerebrovascular injury and its sequelae in adult zebrafish. bioRxiv : the preprint server for biology, doi:10.1101/2025.11.14.688311 (2025).

52 Mary, J. & Jagadeeswaran, P. Zebrafish Model for Thrombosis and Brain-Behavior Studies. Current Protocols 5, e70096, 10.1002/cpz1.70096 (2025).

53 Mizoguchi, T. et al. Neurological function is restored post-ischemic stroke in zebrafish, with aging exerting a deleterious effect on its pathology. Brain Research Bulletin 221, 111225, 10.1016/j.brainresbull.2025.111225 (2025).

54 Porter, B. A. & Mueller, T. The Zebrafish Amygdaloid Complex – Functional Ground Plan, Molecular Delineation, and Everted Topology. Frontiers in Neuroscience Volume 14 - 2020, doi:10.3389/fnins.2020.00608 (2020).

55 Schmidt, R., Strähle, U. & Scholpp, S. Neurogenesis in zebrafish - from embryo to adult. Neural development 8, 3, doi:10.1186/1749-8104-8-3 (2013).

56 Cheng, R. K., Jesuthasan, S. J. & Penney, T. B. Zebrafish forebrain and temporal conditioning. Philosophical transactions of the Royal Society of London. Series B, Biological sciences 369, 20120462, doi:10.1098/rstb.2012.0462 (2014).

57 Fotowat, H., Lee, C., Jun, J. J. & Maler, L. Neural activity in a hippocampus-like region of the teleost pallium is associated with active sensing and navigation. eLife 8, e44119, doi:10.7554/eLife.44119 (2019).

58 Portavella, M., Torres, B. & Salas, C. Avoidance response in goldfish: emotional and temporal involvement of medial and lateral telencephalic pallium. The Journal of neuroscience : the official journal of the Society for Neuroscience 24, 2335–2342, doi:10.1523/jneurosci.4930-03.2004 (2004a).

59 Portavella, M., Torres, B., Salas, C. & Papini, M. R. Lesions of the medial pallium, but not of the lateral pallium, disrupt spaced-trial avoidance learning in goldfish (Carassius auratus). Neuroscience letters 362, 75–78, doi:10.1016/j.neulet.2004.01.083 (2004b).

60 Hasani, H. et al. Whole-brain imaging of freely-moving zebrafish. Frontiers in Neuroscience Volume 17 - 2023, doi:10.3389/fnins.2023.1127574 (2023).

61 Hontani, Y. et al. Deep-Tissue Three-Photon Fluorescence Microscopy in Intact Mouse and Zebrafish Brain. Journal of visualized experiments : JoVE, doi:10.3791/63213 (2022).

62 Murashova, L. & Dyachuk, V. Modeling traumatic brain and neural injuries: insights from zebrafish. Frontiers in molecular neuroscience 18, 1552885, doi:10.3389/fnmol.2025.1552885 (2025).

63 Ochenkowska, K., Herold, A. & Samarut, É. Zebrafish Is a Powerful Tool for Precision Medicine Approaches to Neurological Disorders. Frontiers in molecular neuroscience Volume 15 - 2022, doi:10.3389/fnmol.2022.944693 (2022).

64 Saleem, S. & Kannan, R. R. Zebrafish: A Promising Real-Time Model System for Nanotechnology-Mediated Neurospecific Drug Delivery. Nanoscale research letters 16, 135, doi:10.1186/s11671-021-03592-1 (2021).

65 Shcherbakov, D. et al. Magnetosensation in zebrafish. Current Biology 15, R161–R162, 10.1016/j.cub.2005.02.039 (2005).

66 Shin, J.-N. et al. Zebrafish EEG predicts the efficacy of antiepileptic drugs. Frontiers in Pharmacology Volume 13 - 2022, doi:10.3389/fphar.2022.1055424 (2022).

67 Rahmat, S. & Gilland, E. Hindbrain neurovascular anatomy of adult goldfish (Carassius auratus). Journal of Anatomy 235, 783–793, 10.1111/joa.13026 (2019).

68 Grodzinski, Z. The main vessels of the brain in rainbow trout. Bullet de l’Acad Polonaise des Sciences et des Lettres Serie B: Sci Naturel (II), Juin, 1–19 (1946).

69 Isogai, S., Horiguchi, M. & Weinstein, B. M. The Vascular Anatomy of the Developing Zebrafish: An Atlas of Embryonic and Early Larval Development. Developmental Biology 230, 278–301, 10.1006/dbio.2000.9995 (2001).

70 Lee, N. J. & Matsuoka, R. L. Generation of brain vascular heterogeneity: recent advances from the perspective of angiogenesis. Neural regeneration research 20, 2013–2014, doi:10.4103/nrr.nrr-d-24-00496 (2025).

71 Li, X. et al. A spatiotemporal atlas of cerebrovascular development in zebrafish. Nature Communications 17, 2216, doi:10.1038/s41467-026-68995-z (2026).

72 Gall, L. G. et al. Zebrafish glial-vascular interactions progressively expand over the course of brain development. iScience 28, 111549, doi:10.1016/j.isci.2024.111549 (2025).

73 Panagopoulos, D., Karydakis, P. & Themistocleous, M. The entity of the trapped fourth ventricle: A review of its history, pathophysiology, and treatment options. Brain circulation 7, 147–158, doi:10.4103/bc.bc_30_21 (2021).

74 Ulrich, F., Ma, L.-H., Baker, R. G. & Torres-Vázquez, J. Neurovascular development in the embryonic zebrafish hindbrain. Developmental Biology 357, 134–151, 10.1016/j.ydbio.2011.06.037 (2011).

75 Henderson, J. T. & Piquette-Miller, M. Blood–brain barrier: An impediment to neuropharmaceuticals. Clinical Pharmacology & Therapeutics 97, 308–313, 10.1002/cpt.77 (2015).

76 He, X., Xiong, D., Zhao, L., Fu, J. & Luo, L. Meningeal lymphatic supporting cells govern the formation and maintenance of zebrafish mural lymphatic endothelial cells. Nature Communications 15, 5547, doi:10.1038/s41467-024-49818-5 (2024).

77 Nagata, K., Nakase, H., Kakizaki, T., Otsuka, H. & Sakaki, T. The effect of brain compression under venous circulatory impairment. Neurological Research 22, 713–720, doi:10.1080/01616412.2000.11740745 (2000).

78 Grandel, H., Kaslin, J., Ganz, J., Wenzel, I. & Brand, M. Neural stem cells and neurogenesis in the adult zebrafish brain: Origin, proliferation dynamics, migration and cell fate. Developmental Biology 295, 263–277, 10.1016/j.ydbio.2006.03.040 (2006).

79 Labusch, M., Mancini, L., Morizet, D. & Bally-Cuif, L. Conserved and Divergent Features of Adult Neurogenesis in Zebrafish. Frontiers in Cell and Developmental Biology Volume 8 - 2020, doi:10.3389/fcell.2020.00525 (2020).

80 Obermann, J. et al. The Surface Proteome of Adult Neural Stem Cells in Zebrafish Unveils Long-Range Cell-Cell Connections and Age-Related Changes in Responsiveness to IGF. Stem Cell Reports 12, 258–273, 10.1016/j.stemcr.2018.12.005 (2019).

81 Kishimoto, N., Shimizu, K. & Sawamoto, K. Neuronal regeneration in a zebrafish model of adult brain injury. Disease models & mechanisms 5, 200–209, doi:10.1242/dmm.007336 (2012).

82 März, M., Schmidt, R., Rastegar, S. & Strähle, U. Regenerative response following stab injury in the adult zebrafish telencephalon. Developmental dynamics : an official publication of the American Association of Anatomists 240, 2221–2231, doi:10.1002/dvdy.22710 (2011).

83 Mueller, T. & Wullimann, M. F. An evolutionary interpretation of teleostean forebrain anatomy. Brain, behavior and evolution 74, 30–42, doi:10.1159/000229011 (2009).

84 Westerfield, M. & Zfin. The zebrafish book : a guide for the laboratory use of zebrafish Danio (Brachydanio) rerio. 4th ed. edn, (ZFIN, 2000).

85 Aleström, P. et al. Zebrafish: Housing and husbandry recommendations. Laboratory animals 54, 213–224, doi:10.1177/0023677219869037 (2020).

86 Clark, K. J., Boczek, N. J. & Ekker, S. C. Stressing zebrafish for behavioral genetics. Reviews in the neurosciences 22, 49–62, doi:10.1515/rns.2011.007 (2011).

87 Dahlén, A., Wagle, M., Zarei, M. & Guo, S. Heritable natural variation of light/dark preference in an outbred zebrafish population. Journal of neurogenetics 33, 199–208, doi:10.1080/01677063.2019.1663846 (2019).

88 Monson, C. A. & Sadler, K. C. Inbreeding depression and outbreeding depression are evident in wild-type zebrafish lines. Zebrafish 7, 189–197, doi:10.1089/zeb.2009.0648 (2010).

89 Ayala-Soldado, N., Mora-Medina, R., Molina-López, A. M., Lora-Benítez, A. J. & Moyano-Salvago, R. Evaluation of the Effectiveness of Eugenol and MS-222 as Anesthetics in Zebrafish in Repeated Exposures and Post-Anesthesia Behaviour. Animals : an open access journal from MDPI 14, doi:10.3390/ani14162418 (2024).

90 Collymore, C., Tolwani, A., Lieggi, C. & Rasmussen, S. Efficacy and safety of 5 anesthetics in adult zebrafish (Danio rerio). Journal of the American Association for Laboratory Animal Science : JAALAS 53, 198–203 (2014).

